# In vitro neural networks minimise variational free energy

**DOI:** 10.1101/323550

**Authors:** Takuya Isomura, Karl Friston

**Affiliations:** Laboratory for Neural Computation and Adaptation, RIKEN Center for Brain Science, 2-1 Hirosawa, Wako, Saitama 351-0198, Japan; Wellcome Trust Centre for Neuroimaging, Institute of Neurology, University College London, 12 Queen Square, London, WC1N 3BG, UK

## Abstract

In this work, we address the neuronal encoding problem from a Bayesian perspective. Specifically, we ask whether neuronal responses in an *in vitro* neuronal network are consistent with ideal Bayesian observer responses under the free energy principle. In brief, we stimulated an *in vitro* cortical cell culture with stimulus trains that had a known statistical structure. We then asked whether recorded neuronal responses were consistent with variational message passing (i.e., belief propagation) based upon free energy minimisation (i.e., evidence maximisation). Effectively, this required us to solve two problems: first, we had to formulate the Bayes-optimal encoding of the causes or sources of sensory stimulation, and then show that these idealised responses could account for observed electrophysiological responses. We describe a simulation of an optimal neural network (i.e., the ideal Bayesian neural code) and then consider the mapping from idealised *in silico* responses to recorded *in vitro* responses. Our objective was to find evidence for functional specialisation and segregation in the *in vitro* neural network that reproduced *in silico* learning via free energy minimisation. Finally, we combined the *in vitro* and *in silico* results to characterise learning in terms of trajectories in a variational information plane of accuracy and complexity.

## Introduction

Making inferences about the causes of sensory inputs is one of the most remarkable and essential abilities of animals [1–3]. A famous example of this capability is the cocktail party effect — a partygoer can distinguish an individual′s voice from the noise of a crowd [4,5]. The ability to recognise the cause of particular sensations has been modelled as blind source separation [6–11]. More generally, inferring the (hidden) causes of (observed) sensations constitutes a problem of Bayesian inference [12,13]. In this setting, it is assumed that sensory inputs are generated by mixtures of hidden causes or sources. The aim is then to invert the mapping from causes to consequences and thereby infer the hidden sources — and learn the mapping — using some form of inference. Here, we formalise inference in terms of approximate Bayesian inference; namely the minimisation of variational free energy [13]. This minimisation corresponds to maximising Bayesian model evidence and represents a fundamental form of (unsupervised) learning that may be operating in the brain [14,15].

Interestingly, inference about hidden variables — based on observed data — is a ubiquitous problem in neuroscience: researchers use exactly the same strategy to analyse neuronal (and behavioural) data. Common examples here are general linear models (GLM) and dynamic causal models (DCM) of functional magnetic resonance imaging (fMRI) and electrophysiological time series [16]. These forward or generative models suppose that the signals are generated by hidden (neuronal) dynamics. The inversion of these generative models allows one to infer the hidden variables and learn the model parameters — in the same way that a creature infers the hidden state of its world based on sensory information. In this work, we call on both instances of inference; namely, we try to infer how neurons make inferences. Specifically, we ask how neurons infer the causes of their inputs.

To establish an ideal Bayesian encoding of the hidden causes of sensory stimulation, it is necessary to establish a mapping between idealised (Bayesian) responses (i.e., the sufficient statistics of posterior beliefs about hidden causes) and neuronal responses recorded electrophysiologically. In brain imaging, one would usually use some form of statistical parametric mapping (SPM) or multivariate analysis to identify neuronal populations responding in a way that is consistent with normative principles. Generally, this implies some form of *functional specialisation* and segregation [17,18]. We use the same approach here, to establish a segregation of functionally specialised responses. In other words, we looked for evidence for differential responses to hidden causes of stimuli that emerge, in an experience dependent fashion, over time. However, in our empirical setup, we were not looking at an *in vivo* brain but an *in vitro* neuronal network of cultured cortical cells. These cultures are known to perform various adaptation and learning tasks [19–28] and offer the key advantages of low experimental noise and the opportunity for invasive manipulations. In brief, we analysed *in vitro* neuronal network preparations, which we hypothesised perform Bayesian inference in accordance with the free energy principle [29].

In what follows, we briefly review idealised responses in terms of belief updating under a generative model. We then turn to the analysis of empirical data from cultured cortical cells, looking for evidence of functionally specialised responses due to learning. These empirical responses are then reproduced *in silico* using Bayesian belief updating. Finally, we consider how the synthetic and empirical responses can be combined to characterise real neuronal responses in terms of inference and learning, via free energy minimisation. In short, we tested the hypothesis that *in vitro* neuronal networks show an inherent capacity for self-organising, self-evidencing, free energy minimising behaviour.

## Bayesian inference and learning

### Bayesian source separation by neuronal networks

In our empirical setup, stimuli were generated stochastically by delivering a train of impulses every second to a (8 × 8) grid of stimulation sites of a culturing device (see Methods). The stimuli were generated from two binary sources (*s*_1_ and *s*_2_) and applied probabilistically to two pools of electrodes. One source preferentially excited one pool of electrodes, while the other stimulated a second pool. This stimulation setup speaks to a simple generative process, in which there are two hidden states causing sensory inputs, which can either be active or inactive during each (one second) stimulation epoch. The corresponding likelihood of the implicit generative model can be summarised in a simple likelihood matrix **A**, whose elements correspond to the probability (75% or 25%) of any electrode receiving a stimulus, conditioned on whether the source was present on each trial. With an appropriate generative model, an ideal Bayesian observer should be able to learn and infer the best explanation for this multidimensional sensory input, in terms of the presence or absence of two independent sources.

Intuitively, this sort of process entails changes in synaptic connectivity among the cultured neurons that enables one or more *specialised* neurons to respond selectively to the presence of a particular source. On this view, specialised neurons ‘see’ all inputs vicariously, via connections with other neurons. Crucially, these selective responses emerge despite the fact that no particular input pattern is ever repeated. The emergence of specialised neurons therefore depends upon learning the likelihood of a particular pattern, given the presence of a particular source; i.e., encoding the **A** matrix in terms of synaptic connections. This learning underwrites selective responses that effectively encode the presence or absence of a source. This inference resolves the blind source separation (i.e., the cocktail party) problem formulated, in this instance, in terms of discrete states. To model this source separation, we used a Markov decision process (MDP) model and a biologically plausible gradient descent on variational free energy — as a proxy for log model evidence (i.e., an evidence or marginal likelihood bound). See Fig. 1 for a schematic illustration of how this sort of neuronal (variational) message passing follows from Bayesian inference. For a more complete treatment, please see [30, 31] and Methods.

**Figure 1.**
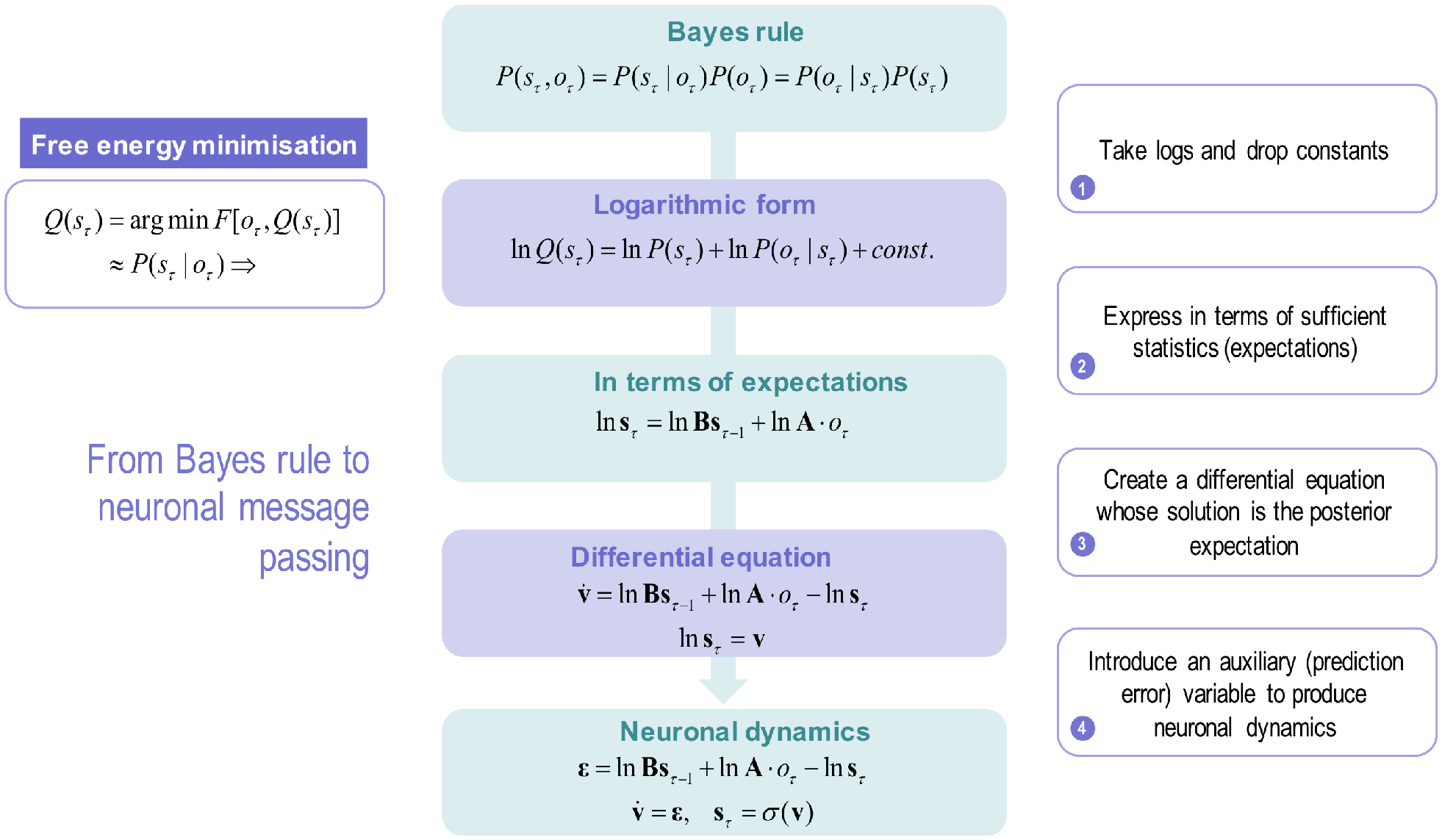
This schematic summarises the conceptual moves that provide a neuronal process theory for Bayesian belief updating or propagation with neuronal dynamics (please see Methods for a more technical and complete description). First, we start with Bayes rule, which says that the joint probability of some causes (hidden states in the world: *s*_τ_) and their consequences (observable outcomes: *o*_τ_) is the same as the probability of causes given outcomes times the probability of outcomes, which is the same as the probability of the outcomes given their causes times the probability of the causes: i.e., *P*(*s*_τ_,*o*_τ_) = *P*(*s*_τ_|*o*_τ_)*P*(*o*_τ_) = *P*(*o*_τ_|*s*_τ_)*P*(*s*_τ_). The second step involves taking the logarithm of these probabilistic relationships and dispensing with the probability over outcomes (because it does not change with the posterior probability of the hidden states we want to infer). Note, at this point, we have replaced the posterior probability with its approximate, free energy minimising, form: *Q*(*s*_τ_) ≈ *P*(*s*_τ_|*o*_τ_). The second step involves rewriting the logarithmic form in terms of the sufficient statistics or parameters of the probability distributions. For discrete state-space models, these are simply expectations (denoted by boldface). Here, we have used an empirical prior; namely, the probability of the current state given the previous state of affairs. The probability transition matrix — entailed by this empirical prior — is denoted by **B**, while the likelihood matrix is denoted by **A**. The fourth move is to write down a differential equation, whose solution is the posterior expectation in the middle panel (expressed as a log expectation). Effectively, this involves introducing a new variable that we will associate with voltage or depolarisation **v**_τ_, which corresponds to the log expectation of causes (**s**_τ_). Finally, we introduce an auxiliary variable called prediction error **ε**_τ_ that is simply the difference between the current log posterior and the prior and likelihood messages. This can be associated with presynaptic drive (from error units) that changes transmembrane potential or voltage (in principal cells); such that the posterior expectation we require is a sigmoid (i.e., activation) function of depolarisation. In other words, expectations can be treated as neuronal firing rate. In summary, starting from Bayes rule and applying a series of simple transformations, we arrive at a neuronally plausible set of differential equations that can be interpreted in terms of neuronal dynamics.

### The generative (Markov decision process) model

Generative models of discrete outcomes are usually described in terms of a likelihood **A** matrix mapping from hidden states of the world and outcomes or sensory input. In addition, they are usually equipped with a probability transition matrix that affords empirical priors on the dynamics of the generating or source process. In the setup described in this paper, the dynamics were very simple. This allows us to focus on inference about the sources currently responsible for generating sensory stimulation. In subsequent work, we will use exactly the same formalism to model sequential stimuli with structured transition probabilities; however, here we consider only one time step for each trial, which means we can ignore the transition probabilities **B**.

The hidden states of this model correspond to a factor (with two levels: present or absent) for every source. In this case, there were two sources leading to (2 × 2 =) 4 hidden states. Given the state of the world delivering stimuli (i.e., the stimulation settings), the elements of the likelihood matrix now specify the probability that any particular electrode would receive an input. These were initially set to a low confidence prior using a Dirichlet parameterisation and initial (concentration) counts of one (i.e., as if the network had only seen one instance of an outcome). A standard Bayesian belief update scheme — variational message passing — was used to update posterior beliefs about sources over successive epochs (see Figs. 1 and 2) and the likelihood a source would generate a stimulus. In other words, the simulated neural network *learned* the likelihood mapping and *inferred* the presence or absence of sources in an experience-dependent fashion. In general, posterior beliefs are encoded by the *sufficient statistics* or parameters of approximate posterior distributions that are associated with neuronal activity and connectivity. Neurobiology plausible process theories of this kind of evidence accumulation mean that we can treat the sufficient statistics (i.e., posterior expectations) about the presence or absence of sources as an idealised neuronal response, engendered by experience-dependent plasticity. Please see [30,32] for details about the belief updating and learning respectively that was used in this paper.

**Figure 2.**
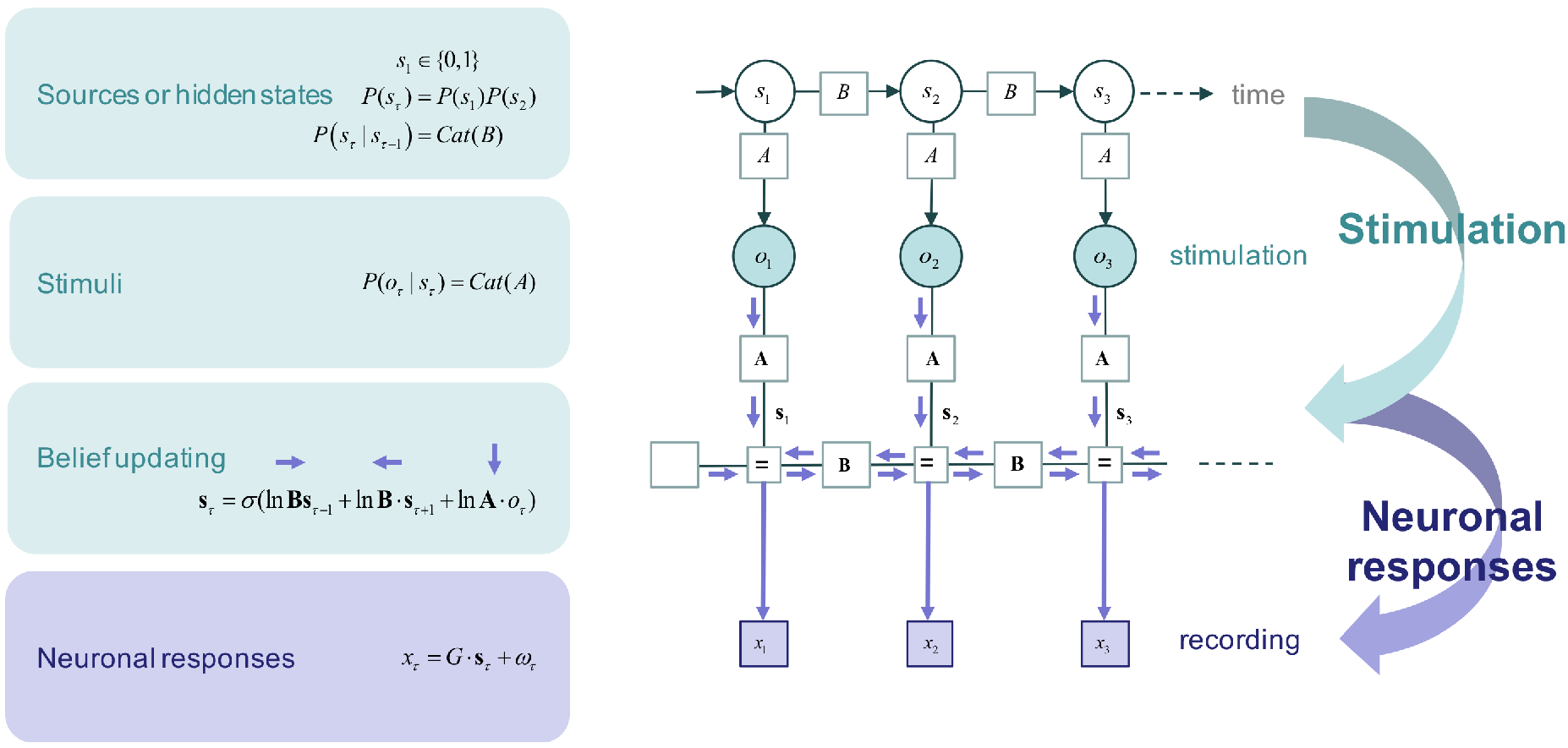
This figure illustrates the belief propagation we used to simulate idealised neuronal responses. This sort of scheme optimises sufficient statistics that encode posterior beliefs about the hidden causes of sensory data. The upper part of this figure uses a graphical model to illustrate how stimuli are generated, while the lower parts of this figure illustrates variational message passing within a neural network — using a Forney factor graph description [55,56] based upon the formulation in [31]. In our setup, we know the hidden states generating observed stimuli — and we have empirical recordings of the sufficient statistics that encode beliefs (or expectations) about these hidden states.

During evidence accumulation over stimulation epochs, the parameters of the likelihood model are accumulated (as Dirichlet concentration parameters); thereby enabling the model to learn which electrodes were likely to be excited by the sources that they were implicitly inferring — and therefore increase the accuracy of posterior beliefs about the sources currently in play.

In this sort of scheme, within-epoch responses are due to a fast gradient descent on variational free energy that combines sensory evidence with prior beliefs entailed by the form of the generative model (see Fig. 2). The between-epoch dynamics correspond to learning the probability with which hidden sources excite any particular electrode. In short, one can simulate ideal responses in terms of the sufficient statistics of the posterior beliefs about the current stimulation pattern (expectations about hidden states) and the contingencies of stimulation (posterior expectations about the likelihood parameter). The key question now is whether these simulated responses — and accompanying decreases in free energy — are evident in empirical neuronal responses.

## Results

### Overview

The empirical responses were summarised in terms of the electrode responses that showed the most significant functional specialisation. We assumed that the deviation of neuronal firing rates from their mean activity is proportional to the difference between the posterior and (uninformative or flat) prior expectations about each of the sources. This is a (thermodynamically) efficient form of neural code because any neuron or population that has not learnt to recognise a hidden state or source will not deviate from prior expectations — and can have a mean firing rate of zero.

Having identified functionally selective empirical responses, we simulated exactly the same sort of recognition process or learning *in silico,* using the above free energy minimising scheme (see Fig. 2). This allowed us to track learning in terms of (simulated) changes in connectivity — using the same stimuli as in the empirical study. These connection strengths correspond to the parameters of the likelihood model (i.e., the **A** matrix) and enabled us to track the accuracy and complexity of inferences about sources based upon the empirical responses (see below).

In what follows, we first describe the emergence of functional specialisation in the empirical data. We then reproduce this learning *in silico.* By matching the time courses of learning, we used the parameters of the generative model from the simulations to assess the empirical free energy associated with neuronal responses (summarised by the most significant electrode). We were hoping to see a progressive decrease in free energy. Furthermore, we were able to decompose the free energy into its constituent parts; namely, *accuracy* and *complexity.* We anticipated that accuracy would increase as learning proceeded, enabling an increase in the complexity or confidence term. To visualise this, we borrowed a construct from the information bottleneck approach; namely an information plane [33–35].

### Estimation of posterior belief based on empirical responses

We first illustrate the emergence of functional specialisation in an exemplar dataset (Fig. 3). These data were obtained over 100 sessions, each of which included 256 stimulation epochs with 1-s intervals based on the above protocol. We used a general linear model (GLM) to characterise the responses observed at each electrode. This GLM comprised explanatory variables (i.e., regressors) using the known sources active in each trial to model the emergence of a differential response to one or the other source. This provides an unbiased estimate of the neuronal encoding of each source after removing influences from the other source and electrode stimulations (i.e., sensory stimulation, from the point of view of the neuron). This modelling ensures that functional specialisation cannot be explained by responses evoked by stimulation *per se* (that are, on average, greater for one source than another). The left panel in Fig. 3A shows the emergence of functional specialisation in terms of the responses at the most significant electrode. The neurons sampled by this electrode become progressively more sensitive to the presence of the first source. To illustrate response selectivity, the responses are shown in red for epochs wherein the first source was present and in cyan for epochs when it was absent.

**Figure 3:**
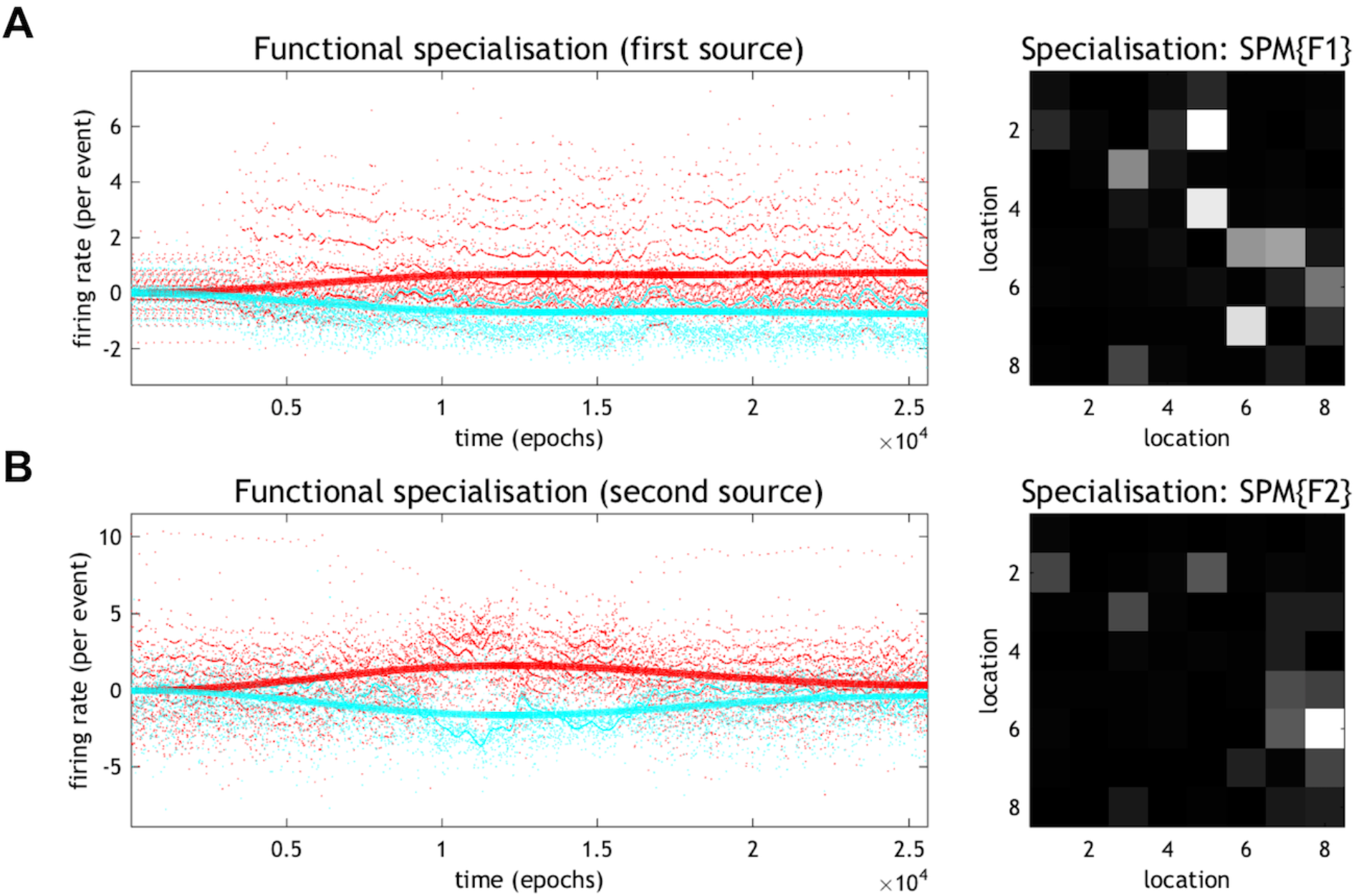
Estimation of blind source separation of functional specialisation from empirical responses. **(A)** Left: The emergence of functional specialisation at the most significant electrode, which became sensitive to the presence of the first source. The red dots correspond to epoch-specific responses when the first source is present, while the cyan dots show the response in the absence of the first source. The red and cyan lines represent the predicted responses; namely, the response associated with the explanatory variables after removal of the effects of stimulation and time. Right: The underlying functional segregation as a statistical parametric map (SPM) of the *F* statistic. This underscores the spatial segregation of functionally specialised responses, when testing for the emergence of selectivity (treating stimulation, non-specific fluctuations, and the other source as confounding effects). The colour scale is arbitrary. Lighter grey colour denotes a more significant effect. **(B)** The analysis presented in these panels is exactly the same as that shown in (A); however, in this case, the explanatory variables modelled an emerging selectivity for the second source.

The explanatory variables in the GLM comprised the interaction between time and the presence of the first source, where time was modelled with a mixture of temporal basis functions (based on a discrete cosine transform with eight components). The best mixture corresponds to the predicted responses shown as solid lines in Fig. 3A. These predicted responses exclude confounding or nuisance effects; namely, the stimulation delivered to the *in vitro* neural network and a discrete cosine transform with 32 components (modelling non-specific drifts in average activity). In short, this GLM enabled the separation of source-specific responses from responses induced by stimulation and fluctuations in the mean response over time. We also observed that the pattern of functional specialisation was distributed but dominated by a small number of electrodes (the right panel in Fig. 3A).

The same analysis was applied to quantify the functional specialisation for the second source and the predicted specialisation is shown in Fig. 3B. This suggests that selectivity in the neurons surrounding these electrodes — for the first and second sources — was circumscribed. One might imagine that other parts of the *in vitro* neural network may have shown selective responses to other patterns of stimuli, or may have indeed shown secondary learning effects at different rates.

The time course of functional specialisation was robust to the choice of neuronal data features. Using the results from 23 cultures, we compared the time course of specialisation obtained using the above procedure (Fig. 4A) with those obtained using alternative data features (Figs. 4B and 4C) and found that the results were very similar. In Fig. 4B, we used the average response over electrodes, whose *F* value exceeded a threshold of 80. In Fig. 4C, the empirical responses were summarised in terms of the principal canonical variate that best explained the emergence of selective responses to each source, in terms of a linear mixture of firing rates. This provided more a sensitive analysis than the corresponding univariate analysis. The canonical variates analysis (CVA) uses the same general linear model that replaces the responses at each electrode with a multivariate response over all electrodes. Finally, specialisation was then compared to the estimates obtained using randomised (surrogate) source trains (Figs. 4D–4F). This nonparametric (null) analysis ensures that the emergence of specialised responses cannot be explained by any confounds or correlations in the data. In summary, we observed the emergence of functional specialisation for both hidden sources, which indicates that *in vitro* neuronal networks can perform blind source separation. Furthermore, this specialisation is robust to the choice of data features and is not an artefact of parametric modelling assumptions.

**Figure 4.**
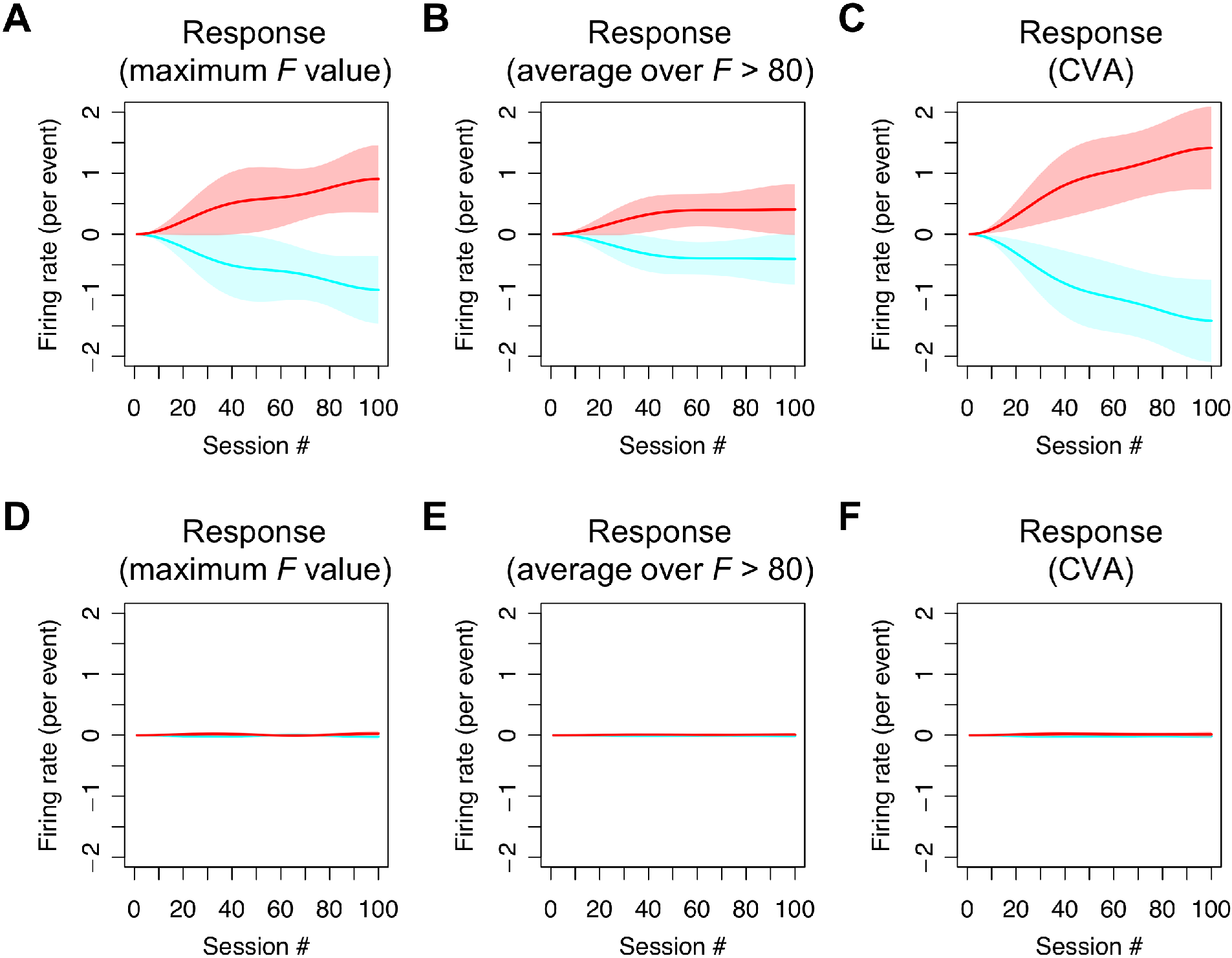
Average responses over culture populations. This figure illustrates the statistical robustness of the procedures used to estimate functional specialisation. Each panel shows the mean differential responses in the presence (red line) and absence (cyan) of the first source, for each session averaged over 23 samples (i.e., cultures). The shaded area indicates the standard deviation over samples. **(A)** Responses obtained based on activity at an electrode with the maximum *F* value, corresponding to the (single culture) result in Fig. 3A. **(B)** Responses obtained using a within-culture average over unit activities at electrodes whose *F* value exceeded 80. Here, we first calculated an average response over electrodes within a culture, and then calculated an average over cultures. **(C)** Responses obtained using a standard canonical variate analysis (CVA) with the GLM described in the main text. This shows the emergence of functional specialisation in terms of the first canonical variate (i.e. pattern over electrodes) that becomes specialised for the first source. This canonical variate represents a linear mixture of firing rates from all electrodes, averaged over each epoch. Panels **(D)**, **(E)**, and **(F)** are the same as panels **(A)**, **(B)**, and **(C)**, but using surrogate (randomised) source signals. The comparison of the upper and lower panels illustrates the emergence of functional specialisation when, and only when, the true sources were used.

### Simulation of belief updates and learning in the Bayes optimal encoder

To establish that the functional specialisation observed above conforms to Bayesian learning (i.e., the free energy principle), we simulated learning under ideal Bayesian assumptions (Fig. 5). This scheme corresponds to evidence accumulation under the hidden Markov model (described in Fig. 2) with state transitions **B** modelled by an identity matrix. The hidden states were inferred based on the same stimuli used in the empirical experiment. For computational expediency, we simulated learning over 512 epochs. Figure 5 illustrates the emergence of specialised responses in terms of firing rates encoding the presence and absence of the first source (Fig. 5A), the time frequency profile of induced responses (Fig. 5B) and underlying local field potential (Fig. 5C). These simulated neurophysiological responses are based on the process theory summarised in Fig. 1 (and explained in detail in [30]). The resulting emergence of selective responses (i.e., learning) is shown in Fig. 5D using the same GLM analysis and format of Fig. 3; namely, the analysis of the empirical data. Similar results were obtained for the units encoding the presence of the other source. The correspondence between the empirical and synthetic responses is self-evident.

**Figure 5.**
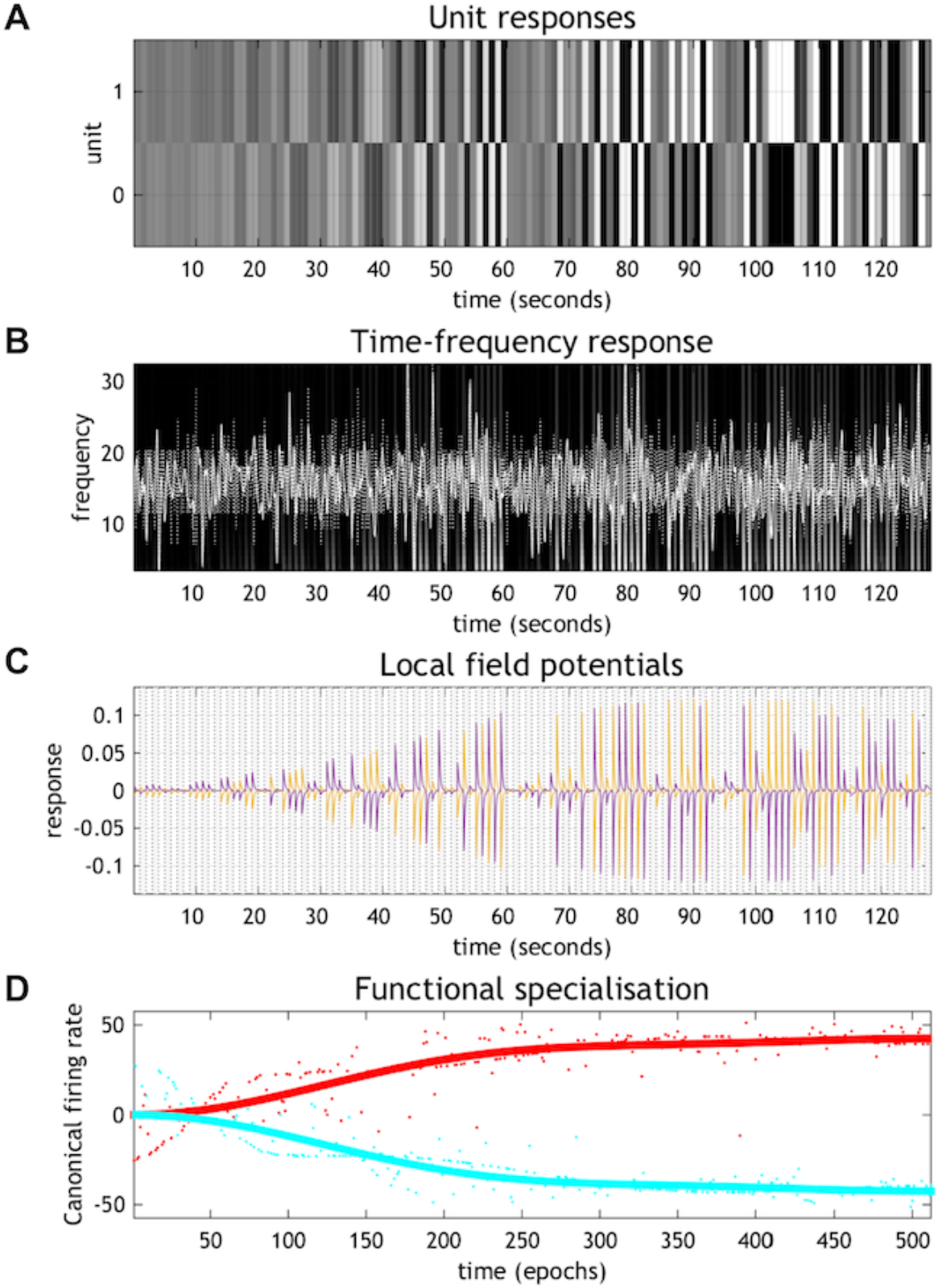
Synthetic responses of the simulated Bayes optimal encoder. **(A)** Simulated firing rates for the first 128 epochs, focusing on units encoding the absence (top) and presence (bottom) of the first source. **(B)** The equivalent responses averaged over all neurons after band-pass filtering (white lines). These simulated local field potentials are shown on a background image of induced responses following a time frequency analysis (see [30] for details). **(C)** A more detailed representation of the simulated local field potentials of the units shown in panel **(A)**. These field potentials are the band-pass filtered firing rates of the unit encoding the posterior expectation of one source (purple line) and its absence (yellow line). **(D)** The resulting emergence of selective responses, plotted in the same format used in Fig. 3, where red and cyan lines express responses in the presence and absence of the first source, respectively.

The above analyses show that Bayes optimal encoding of the causes of sensory stimulation provides a qualitative account of observed electrophysiological responses. In what follows, we quantify the empirical responses in terms of the inference and learning, by matching the time course of synthetic and empirical learning. This allowed us to interpret the empirical responses in terms of posterior beliefs about hidden sources.

### Integrating empirical and synthetic results

In this final section, we quantify the progressive reduction in variational free energy, during the emergence of functional specialisation in the cell culture (Fig. 6). This characterisation combines the results presented in the previous two sections to evaluate the free energy (and its components) entailed by the empirical responses. In detail, the time course of learning was quantified using the appropriate mixture of temporal basis functions from the analyses of empirical and synthetic responses. The corresponding learning curves were matched, in a least squares sense, by regressing the empirical learning on the synthetic learning curves (see Fig. 6D). For a selected series of 512 equally spaced empirical epochs, the corresponding point in learning was identified in the simulations. The parameters of the generative model **A**_*t*_ were then used to evaluate the variational free energy using the stimulation pattern *o*_*t*_ and the empirical posterior expectations: **s**_*t*_. These empirical expectations were derived in a straightforward way by assuming that the predicted specialisation (i.e., red lines in Fig. 3) corresponded to a posterior expectation of one. With these expectations, we can now evaluate the empirical free energy for each epoch *t*: defined as follows (please see Methods for details):

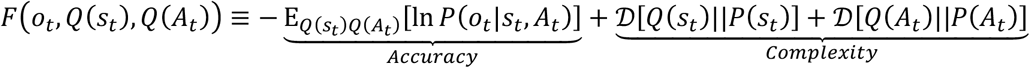

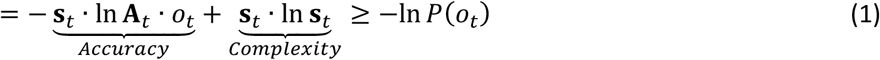

Here, *Q*(*s*_*t*_) and *Q*(*A*_*t*_) denote approximate posterior distributions over hidden states encoded by neuronal activity and elements of the likelihood matrix encoded by synaptic connectivity, respectively. The free energy has been expressed here in terms of *accuracy* and *complexity;* namely, the expected log likelihood of sensory outcomes and the Kullback-Leibler divergence between posterior and prior beliefs. The inequality indicates that variational free energy is an upper bound on negative log evidence or marginal likelihood *P*(*o*_*t*_) of sensory outcomes at any particular time. Note that in the second line, complexity of matrix **A** is omitted because it is determined by the simulation and does not change the results in this section.

**Figure 6.**
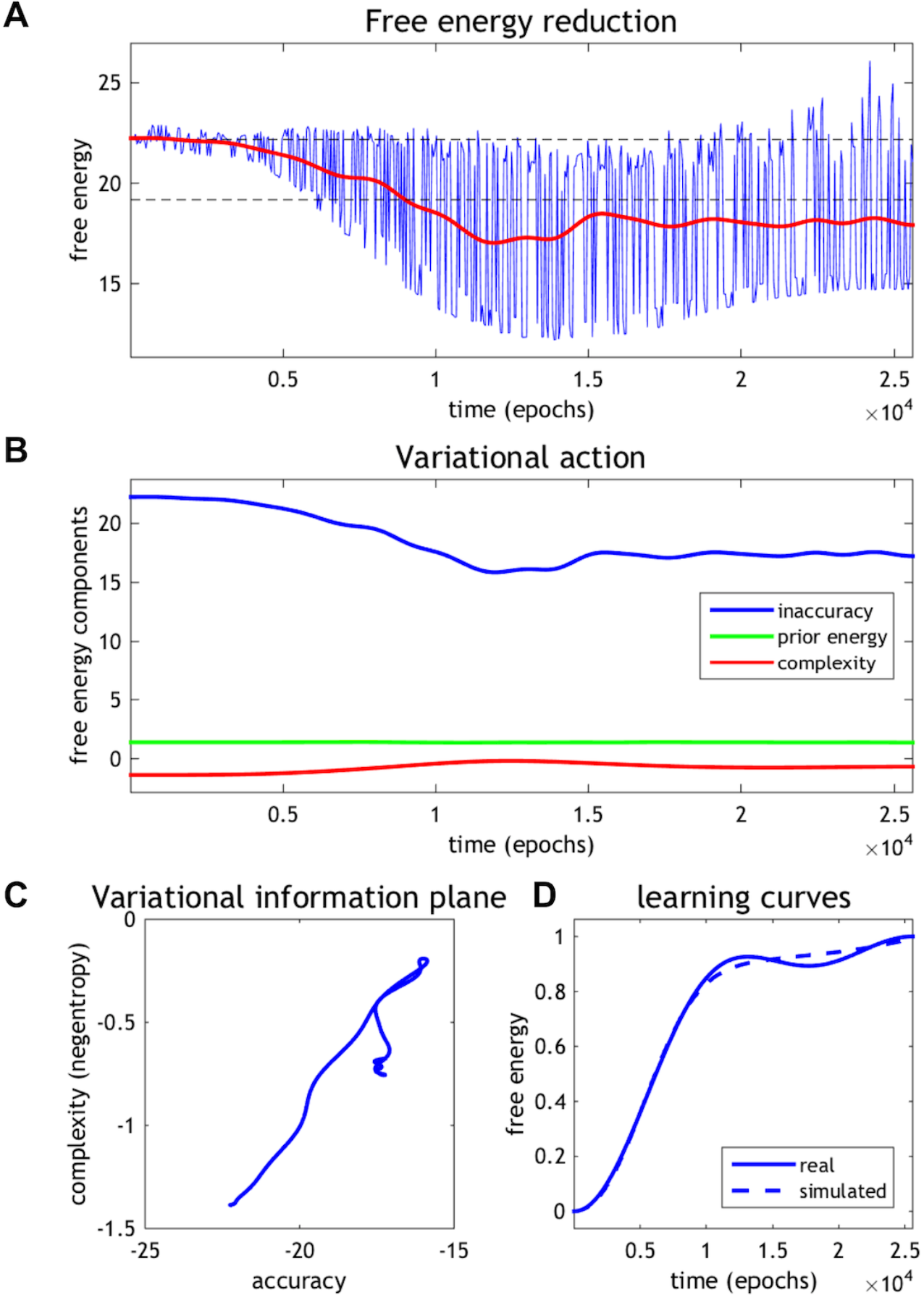
Empirical free energy minimisation. **(A)** The resulting fluctuations in free energy. The blue line corresponds to the free energy based upon the neuronal encoding (the lines in Fig. 3) and the red line shows the average over 32 successive epochs. **(B)** Trajectories of free energy components after smoothing. **(C)** Variational information plane. **(D)** Trajectories of learning curves obtained from empirical (solid line) and simulated (dashed line) data.

The resulting fluctuations in free energy are shown in Fig. 6A. The blue line corresponds to the free energy based upon the predicted neuronal encoding (i.e., calculated based on data in Fig. 3 using the above equation). To make the emergence of functional specialisation or learning easier to visualise, we plotted the same data after the free energy was smoothed using a time window of 32 epochs (the red line in Fig. 6A). This average suppresses epoch to epoch fluctuations due to the different patterns of stimulation that the neurons find more or less easy to recognise — in terms of the sources that caused them. The horizontal lines show the free energy at the beginning of the session and the same value after subtracting three nats (i.e., natural units). This allows a quantitative interpretation of the free energy. This is because a free energy difference of about three corresponds to strong evidence for the presence of a source; i.e., a log odds ratio of exp(3) = 20 to 1. Note that by the end of the session, nearly every pattern of stimulation is recognised in terms of its underlying sources, in virtue of having a relatively low free energy, or high model evidence.

Figure 6B decomposes the components of free energy into the negative log likelihood (*inaccuracy*), negative log prior, and negative entropy (*complexity*). It can be seen that the accuracy (i.e., the log likelihood) increases dramatically as learning proceeds — and evidence has accumulated about the causes of stimulation. This is accompanied by a smaller increase in complexity or a decrease in entropy, as inferences about the sources become more confident. We anticipated that accuracy would increase as learning proceeded, enabling an increase in the complexity or confidence term. To visualise this, we borrowed a construct from the information bottleneck approach; namely, an information plane [30]. However, in this application we plot accuracy against complexity in a *variational information plane*. Note that, under flat or uninformative priors, complexity corresponds to the negative entropy of free energy (i.e., negentropy). This means we can associate negative accuracy (inaccuracy) with energy and complexity with negentropy.

Figure 6C shows the evolution of accuracy and complexity in a variational information plane as functional specialisation emerges. This shows that as learning progresses, accuracy increases markedly with an accompanying increase in complexity. However, there is an interesting exception to this trend; it appears as if the complexity falls later training period. Anecdotally, the network appears to first maximise accuracy (with an accompanying complexity cost) and then tries to find a less complex explanation. This is reminiscent of the behaviour of deep learning schemes described in [33–35]. We confirmed that this tendency was conserved over 23 recording samples (Fig. 7A). In almost half of the cultures, complexity falls after an initial increase. Complexity, in this context, corresponds to the divergence between posterior and prior beliefs and can be thought of as the degrees of freedom used in providing an accurate explanation for observed data. In summary, while the accuracy increased progressively over the training period, the complexity increased initially and then either levelled off or fell. This is consistent with Ockham’s (and the free energy) principle, in the sense that this behaviour can be interpreted as trying to find a simpler explanation for observed outcomes. Moreover, this tendency was robust to the choice of data features (Figs. 7B and 7C). Finally, these systematic trajectories in the information plane disappeared when decomposing the free energy obtained using randomised (surrogate) source trains (Fig. 7D).

**Figure 7.**
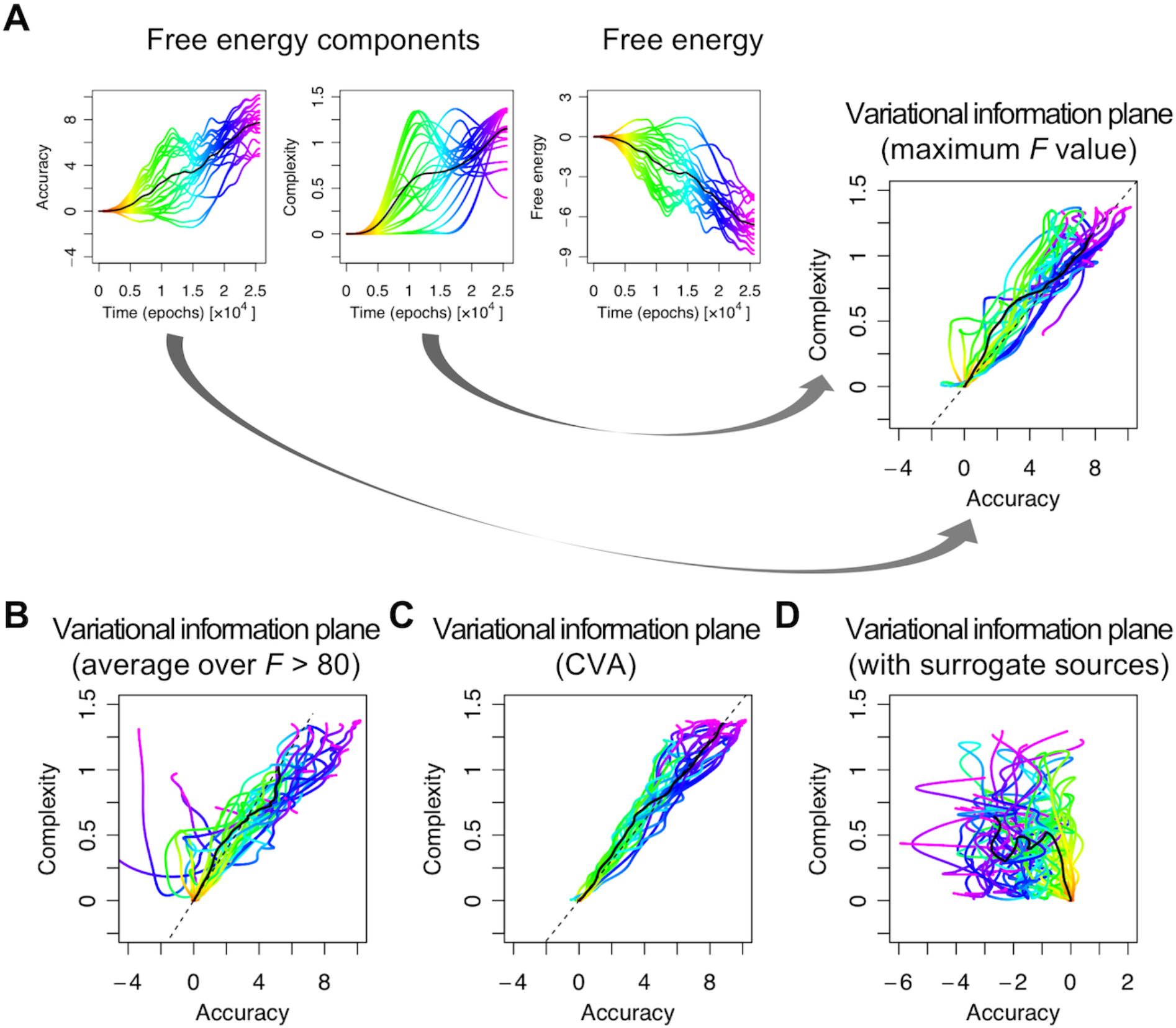
Variational information plane analysis. **(A)** Trajectories of accuracy, complexity (negentropy), and free energy as a function of time or learning. These trajectories were based on the responses at the electrode with the maximum value of the *F* statistic. Each coloured line corresponds to a different culture. The colour indicates the time course. The black lines report the average over 23 cultures. The rightmost panel shows the corresponding trajectories in the information plane by plotting complexity against accuracy. Panels **(B)** and **(C)** have the same format as panel **(A)**, but different data features were used to evaluate free energy; namely, responses obtained using a within-culture average over electrodes whose *F* value exceeded a threshold **(B)** or responses obtained using a canonical variate analysis **(C)**. Panel **(D)** is the same as the information plane in panel **(A)**, but using surrogate (i.e. randomise) sources. These null results suggest that accuracy actually fell over time to a small extent with learning.

## Discussion

In this study, we have demonstrated that *in vitro* neuronal cell cultures can recognise and learn statistical regularities in the patterns of their stimulation — to the extent they can be regarded as performing blind source separation or perceptual inference on the causes of their input. According to normative variational principles for the brain [12,36–38], neural encoding entails the inference and representation of information in the external world; i.e., the hidden sources or causes of sensory consequences. We therefore tried to infer the neural code that underwrites this representation. Formally, this is a meta-Bayesian problem; in the sense that we are trying to make inferences about empirical neuronal recordings that are generated by a process of Bayesian inference. We simulated an ideal Bayesian neural network and then considered the mapping from idealised responses (i.e., the ideal neural code) to recorded neuronal activity. We found clear evidence for learning and inference via the emergence of functional specialisation in the empirical data. Furthermore, this specialisation was robust to the choice of neuronal data features and mirrored simulated (idealised) specialisation. By establishing a mapping of the *in vitro* and *in silico* responses, we were able to evaluate the posterior ′beliefs′ about hidden sources associated with empirical responses — and demonstrate a significant reduction in free energy over the course of the training.

We adopted a generative model of discrete sensory outcomes described in terms of a likelihood **A** matrix, mapping hidden causes in the external world to the observable outcomes, as in the cocktail party problem [4,5] and blind source separation [6–11]. This simple setup allowed us to characterise neuronal responses encoding the sources responsible for generating sensory stimulation. Usually these (hidden Markov or Markov process) models are equipped with a probability transition matrix **B** that corresponds to empirical priors on structured sequences. In subsequent work, we will use the same formalism introduced in this paper to model sequential stimuli with structured transition probabilities. In principle, this should provide a full meta-Bayesian approach in which the neuronal encoding model is itself inverted using Bayesian procedures.

Neurobiologically, our Bayesian inference and learning processes describe neuronal activity and synaptic plasticity, respectively (see Eqs. (7) and (8) in Methods). In this setting, the softmax function used to evaluate the posterior expectation of hidden states, given its logarithmic form, might correspond to a nonlinear activation (i.e., voltage-firing) function. It is well known that the mean firing rate function of spiking neuron models has the analytic form of a softmax function [39]. Moreover, our learning rule corresponds exactly to a Hebbian rule of associative plasticity, where observations and the posterior-expectation-coding neurons correspond to pre and postsynaptic neurons, respectively. This sort of Hebbian plasticity is physiologically implemented as spike-timing dependent plasticity (STDP) [40–43]. These form observations speak to the biological plausibility of the variational message passing scheme used to simulate neural responses in this paper. Although several biologically plausible BSS methods in the continuous state space have been developed [44–48], to our knowledge, this is the first attempt to explain neuronal BSS using a biologically plausible learning algorithm in the discrete (binary) state space.

Furthermore, our results suggest that *in vitro* neuronal networks can perform Bayesian inference and learning under the sorts of generative models assumed in our Markov decision process (MDP) simulations. It is interesting to consider how real neurons actually encode information; for example, synaptic plasticity (i.e., learning) is modulated by the various neurotransmitters (such as GABA) and neuromodulators (such as dopamine and noradrenaline) [49–51]; please see also [52] for a possible relationship between neuromodulations of STDP and the free-energy principle. This implies the existence of a mapping between the variables in the MDP scheme and the concentrations of these neurotransmitters. In the subsequent work, we will touch on this issue by asking how alterations in the level of neurotransmitters and neuromodulators influence Bayesian inference and learning evinced by *in vitro* neural networks.

In summary, we have characterised the neural code in terms of (approximate) Bayesian inference by mapping empirical neuronal responses to inference about the hidden causes of observed sensory inputs. Using this scheme, we were able to demonstrate meaningful reductions in variational free energy in *in vitro* neural networks. Moreover, we observed that the ensuing functional specialisation show some characteristics that are consistent with Ockham’s principle; namely, a progressive increase in accuracy, with an accompanying complexity cost, followed by a simplification of the encoding — and subsequent reduction in complexity. This is similar to a phenomenon observed in a recent deep learning study [33–35]. These results highlight the utility of inference as a basis for understanding the neural code and the function of neural networks.

## Methods

### Cell culture

The dataset used for this study was originally used in the previous study of neuronal blind source separation, and detailed methods can be found in [29]. All spike number data can be downloaded from http://neuron.t.u-tokyo.ac.jp. All animal experiments were performed with the approval of the animal experiment ethics committee at the University of Tokyo (approval number C-12-02, KA-14-2) and according to the University of Tokyo guidelines for the care and use of laboratory animals.

Briefly, the cerebral cortex of 19-day-old embryos (E19) was obtained from pregnant Wistar rats (Charles River Laboratories, Japan) and dissociated into single cells by treatment with 2.5% Trypsin (Life Technologies) followed by mechanical pipetting. A half million (5 × 10^5^) dissociated cortical cells (a mixture of neurons and glial cells) were seeded on the centre of micro electrode array (MEA) dishes and cultivated in the CO_2_ incubator. We used data from 23 cultures for analysis. See [19,53] for the detail about MEA.

### Electrophysiology

Electrophysiological experiments were conducted using an MEA system (NF Corporation, Japan). An MEA dish comprises 8×8 microelectrodes embedded on its centre, deployed on a grid with 250-μm microelectrodes separation. These microelectrodes are dual-use for recording and stimulation, enabling extracellular recordings of evoked spikes (early response) from multiple sites immediately following electrical stimulations. 14-hour recordings were acquired at a 25 kHz sampling frequency and band-pass filtered between 100–2000 Hz. A biphasic pulse of amplitude 1V and 0.2 ms duration was used as a stimulation input. This stimulation pulse is known to efficiently induce activity-dependent synaptic plasticity [19,29]. All recordings and stimulation were conducted in a CO_2_ incubator.

***Generative process:*** We prepared two sequences of hidden sources and applied their stochastic mixtures to neural networks over 32 electrodes. Half (16) of the electrodes were stimulated under source 1, with a probability of 3/4 or source 2, with a probability of 1/4. Conversely, the remaining (16) electrodes were stimulated under source 1, with a probability of 1/4 or source 2, with a probability of 3/4. The 32 stimulated electrodes were randomly selected in advance and fixed over training.

In terms of the MDP scheme [30], this corresponds to the likelihood mapping **A** from two hidden sources or states *s* = *s*_1_⊗*s*_2_ to 32 observations *o* = *o*_1_⊗…⊗*o*_32_ (see also Fig. 2). Each source and observation takes values of zero or one (*s*_1_,*s*_2_ ∈ {0,1}, *o*_*i*_ ∈ {0,1}) for each trial. The probability of *s* follows a uniform (categorical) distribution *P*(*s*) = *P*(*s*_1_)*P*(*s*_2_) = 0.5 × 0.5, while the probability of *o*_*i*_ is determined by a categorical distribution *P*(*o*_*i*_|*s*_1_⊗*s*_2_,*A*) = *Cat*(*A*_*i*_) where each element of **A** is given by *P*(*o*_*i*_ = *j*|*s*_1_ = *k*,*s*_2_ = *l*, *A*) = *A*_*ijkl*_. For the process generating outcomes, this means that the values of the likelihood matrix are:

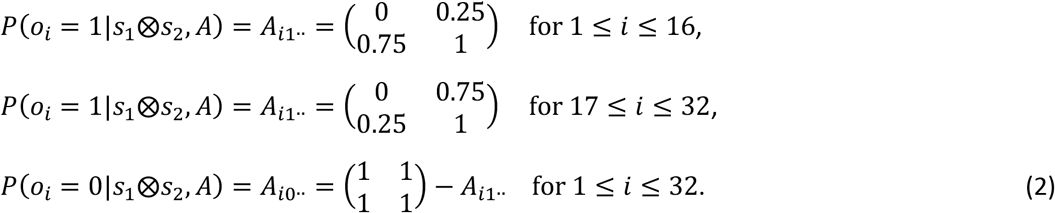

A session of training comprising 256 trials with the 1 s intervals, followed by 244 second rest periods. We repeated this training 500 second cycle for 100 sessions. By inducing electrical stimulation generated by a mixture of hidden sources and monitoring the evoked responses over 14 hours, we were able to characterise the emergence of functional specialisation of certain neurons in these cultured neural networks.

### Analysis

***Estimation of posterior beliefs based on empirical responses:*** The hypothesis we have in mind is that stimulation (*o*) excites a subset of neurons in the tissue culture in an obligatory fashion. With repeated exposure, other neurons — with the right sort of connectivity — will come to recognise the patterns of responses as being caused by the presence or absence of hidden sources (*s*). Following the process theory for this form of Bayesian source separation — based on minimising variational free energy — we assume that the expected probability of sources being present or absent come to be encoded in the firing rate of functionally specialised neurons, whose activity is reported by the recording electrodes.

Under these assumptions, the activity recorded at the 64 electrodes would receive contributions from populations receiving input and neurons encoding the presence of sources. To identify functionally specialised neurons, it is therefore necessary to separate responses directly induced by stimulation from those encoding the sources. In brief, we identified specialised responses by modelling recorded activity as a mixture of stimulation related responses and functionally specialised responses to the sources. Crucially, the former should remain invariant over time, while the latter will emerge during learning. This distinction allows us to decompose responses into stimulation and source specific components — and thereby identify electrodes in functionally segregated regions of the culture.

The time course of this emerging specialisation was modelled using temporal basis functions to avoid any bias in estimating the associated learning rate. Then, the empirical responses were summarised in terms of the electrode responses with the most significant functional specialisation; i.e., a patterns of firing that progressively differentiated between the presence and absence of one of the two sources. In Fig. 3, significant specialisation was identified using empirical responses at electrodes with the greatest *F* statistic. For comparison, in Fig. 4, we used two additional data features. Surrogate analyses were conducted for all analyses to verify they were robust to any (parametric) assumptions.

***Simulated Markov decision process scheme:*** We used the same generative process described above and simulated the Bayes optimal encoding for the causes or sources of sensory stimulation. This was done by implemented using variational message passing has coded in **spm_MDP_VB_X** in the open access academic software SPM (http://www.fil.ion.ucl.ac.uk/spm/software/). See [30] for details regarding this MDP scheme. The posterior expectations of the two sources obtained by the MDP scheme were plotted in Fig. 5.

Briefly, in the MDP scheme, we define a generative model probabilistically with:
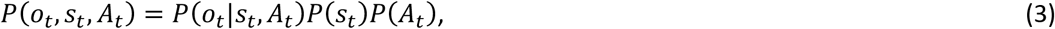

Here, the likelihood is given by a categorical distribution *P*(*o*_*t*_|*s*_*t*_,*A*_*t*_) = *Cat*(*A*_*t*_) and the prior distribution of *A*_*t*_ is given by Dirichlet distribution *P*(*A*_*t*_) = *Dir*(*a*) with sufficient statistics *a*. The mean-field approximation provides an approximation to the posterior (recognition) density:
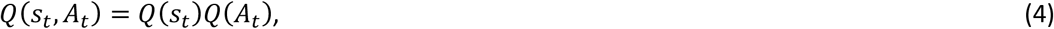

Here, the posterior distributions of *s*_*t*_ and *A*_*t*_ are given by a categorical *Q*(*s*_***t***_) = *Cat*(***s***_***t***_) and Dirichlet *Q*(*A*_*t*_) = *Dir*(***a***_***t***_) distributions, respectively. Note that **s***_t_* and **a**_*t*_ constitute sufficient statistics. Below, we use the posterior expectation of ln*A*_*i,t*_ to express the posterior belief about hidden parameters, which is given by:
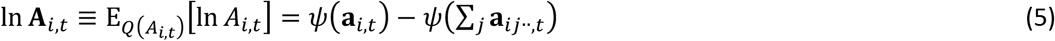

This expression uses the digamma function *ψ*(·). Neurobiologically, this likelihood mapping is associated with synaptic strengths. The free energy of this system is given by:
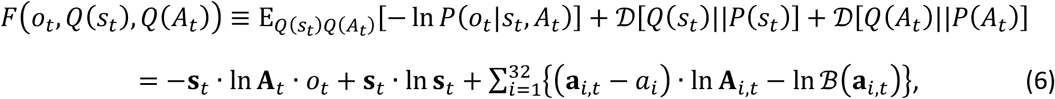

Here, 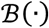 is the beta function. The first term in the right side corresponds to the accuracy, while the second and third terms constitute complexity (expressed by Kullback-Leibler divergence [54]). Inference optimises the posterior expectations of hidden states to minimise free energy, which is given by
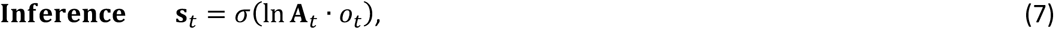

where *σ*(·) is a softmax function (associated with a nonlinear neuronal activation function). While Eq. (7) was derived from the free energy minimisation, the same result can be obtained from Bayes rule as shown in Fig. 1. Moreover, learning entails updating posterior expectations about the parameters to minimise free energy:
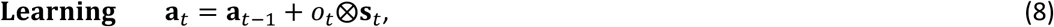

where ⊗ express the operator of outer product. Neurobiologically, Equations (5) and (8) are usually associated with synaptic plasticity (i.e., a Hebbian rule).

***Empirical free energy components:*** We defined **s**_*t*_ as the predicted specialisation obtained in Figs. 3 and 4 and substituted these values into Eq. (6). This enabled us to evaluate the empirical free energy during learning and to decompose free energy into accuracy and complexity, which are shown in Figs. 6 and 7.

The analyses reported in this study represent a point of reference for future studies that will examine the effects of stimulation patterns, sequences of sources and pharmacological manipulations on the learning — as characterised in terms of free energy, accuracy and complexity.

## Acknowledgements

This work was supported by RIKEN Center for Brain Science (T.I.). K.J.F. is funded by a Wellcome Principal Research Fellowship (Ref: 088130/Z/09/Z). The funders had no role in study design, data collection and analysis, decision to publish, or preparation of the manuscript.

## Author Contributions

Conceived and designed the proposed scheme: K.J.F. Performed the simulations: T.I. and K.J.F. Wrote the paper: T.I. and K.J.F.

## Additional Information

### Competing Interests

the authors declare that they have no competing interests.

## References

1 von Helmholtz, H. Treatise on physiological optics (Vol. 3) (The Optical Society of America, 1925).

2 Knill, D. C. & Pouget, A. The Bayesian brain: the role of uncertainty in neural coding and computation. Trends Neurosci. 27, 712–719 (2004).

3 DiCarlo, J. J., Zoccolan, D. & Rust, N. C. How does the brain solve visual object recognition? Neuron 73, 415–434 (2012).

4 Brown, G. D., Yamada, S. & Sejnowski, T. J. Independent component analysis at the neural cocktail party. Trends Neurosci. 24, 54–63 (2001).

5 Mesgarani, N. & Chang, E. F. Selective cortical representation of attended speaker in multi-talker speech perception. Nature 485, 233–236 (2012).

6 Bell, A. J. & Sejnowski, T. J. An information-maximization approach to blind separation and blind deconvolution. Neural Comput. 7, 1129–1159 (1995).

7 Bell, A. J. & Sejnowski, T. J. The “independent components” of natural scenes are edge filters. Vision Res. 37, 3327–3338 (1997).

8 Amari, S. I., Cichocki, A. & Yang, H. H. A new learning algorithm for blind signal separation. Adv Neural Inf Proc Sys. 8, 757–763 (1996).

9 Hyvärinen, A. & Oja, E. One-unit learning rules for independent component analysis. Adv Neural Inf Proc Sys. 9, 480–486 (1997).

10 Cichocki,A., Zdunek, R., Phan, A. H. & Amari, S.I. Nonnegative Matrix and Tensor Factorizations: Applications to Exploratory Multi-way Data Analysis and Blind Source Separation (John Wiley & Sons, 2009).

11 Comon, P. & Jutten, C. Handbook of Blind Source Separation: Independent Component Analysis and Applications (Academic Press, 2010).

12 Friston, K., Kilner, J. & Harrison, L. A free energy principle for the brain. J. Physiol. Paris 100, 70–87 (2006).

13 Friston, K. The free-energy principle: a unified brain theory?. Nat. Rev. Neurosci. 11, 127–138 (2010).

14 Dayan, P. & Abbott, L. F. Theoretical neuroscience: computational and mathematical modeling of neural systems (MIT Press, London, 2001).

15 Gerstner, W. & Kistler, W. Spiking Neuron Models. Single Neurons, Populations, Plasticity (Cambridge University Press, Cambridge, 2002).

16 Friston, K. J., Trujillo-Barreto, N. & Daunizeau, J. DEM: A variational treatment of dynamic systems. NeuroImage 41, 849–885 (2008).

17 Zeki, S. The Ferrier Lecture 1995 behind the seen: the functional specialization of the brain in space and time. Philos. Trans. R. Soc. Lond. B. Biol. Sci. 360, 1145–1183 (2005).

18 Zeki, S.& Shipp, S. The functional logic of cortical connections. Nature 335, 311–317 (1988).

19 Jimbo, Y., Tateno, T. & Robinson, H. P. C. Simultaneous induction of pathway-specific potentiation and depression in networks of cortical neurons. Biophys. J. 76, 670–678 (1999).

20 Chiappalone, M., Massobrio, P. & Martinoia, S. Network plasticity in cortical assemblies. Eur. J.Neurosci. 28, 221–237 (2008).

21 Johnson, H. A., Goel, A. & Buonomano, D. V. Neural dynamics of in vitro cortical networks reflects experienced temporal patterns. Nat. Neurosci. 13, 917–919 (2010).

22 Shahaf, G. & Marom, S. Learning in networks of cortical neurons. J. Neurosci. 21, 8782–8788 (2001).

23 Eytan, D., Brenner, N. & Marom, S. Selective adaptation in networks of cortical neurons. J.Neurosci. 23, 9349–9356 (2003).

24 Le Feber, J., Stegenga, J. & Rutten, W. L. The effect of slow electrical stimuli to achieve learning in cultured networks of rat cortical neurons. PLoS ONE 5, e8871 (2010).

25 Ruaro, M. E., Bonifazi, P. & Torre, V. Toward the neurocomputer: image processing and pattern recognition with neuronal cultures. IEEE Trans. Biomed. Eng. 52, 371–383 (2005).

26 Feinerman, O. & Moses, E. Transport of information along unidimensional layered networks of dissociated hippocampal neurons and implications for rate coding. J. Neurosci. 26, 4526–4534 (2006).

27 Feinerman, O., Rotem, A. & Moses, E. Reliable neuronal logic devices from patterned hippocampal cultures. Nat. Phys. 4, 967–973 (2008).

28 Dranias, M. R., Ju, H., Rajaram, E. & VanDongen, A. M. Short-term memory in networks of dissociated cortical neurons. J. Neurosci. 33, 1940–1953 (2013).

29 Isomura, T., Kotani, K. & Jimbo, Y. Cultured cortical neurons can perform blind source separation according to the free-energy principle. PLoS Comput. Biol. 11, e1004643 (2015).

30 Friston, K., FitzGerald, T., Rigoli, F., Schwartenbeck, P. & Pezzulo, G. Active inference: A process theory. Neural Comput. 29, 1–49 (2017).

31 Friston, K.J., Parr, T. & de Vries, B. D. The graphical brain: belief propagation and active inference. Netw. Neurosci. 1, 381–414 (2017).

32 Friston, K., FitzGerald, T., Rigoli, F., Schwartenbeck, P. & Pezzulo, G. Active inference and learning. Neurosci. Biobehav. Rev. 68, 862–879 (2016).

33 Shwartz-Ziv, R. & Tishby, N. Opening the black box of deep neural networks via information. arXiv preprint arXiv:1703.00810 (2017).

34 Tishby, N. & Zaslavsky, N. Deep learning and the information bottleneck principle. In Information Theory Workshop, 2015, 1–5, IEEE (2015).

35 Saxe, A. M., Bansal, Y., Dapello, J., Advani, M., Kolchinsky, A., Tracey, B. D. & Cox, D. D. On the information bottleneck theory of deep learning. In International Conference on Learning Representations (2018).

36 Dayan, P., Hinton, G. E., Neal, R. M. & Zemel, R. S. The Helmholtz machine. Neural Comput. 7, 889–904 (1995).

37 George, D. & Hawkins, J. Towards a mathematical theory of cortical micro-circuits. PLoS Comput. Biol. 5, e1000532 (2009).

38 Bastos, A. M., Usrey, W. M., Adams, R. A., Mangun, G. R., Fries, P. & Friston, K. J. Canonical microcircuits for predictive coding. Neuron 76, 695–711 (2012).

39 Brunel, N. & Latham, P. E. Firing rate of the noisy quadratic integrate-and-fire neuron. Neural Comput. 15, 2281–2306 (2003).

40 Markram, H., Lübke, J., Frotscher, M. & Sakmann, B. Regulation of synaptic efficacy by coincidence of postsynaptic APs and EPSPs. Science 275, 213–215 (1997).

41 Bi, G. Q. & Poo, M. M. Synaptic modifications in cultured hippocampal neurons: dependence on spike timing, synaptic strength, and postsynaptic cell type. J. Neurosci. 18, 10464–10472 (1998).

42 Froemke, R. C. & Dan, Y. Spike-timing-dependent synaptic modification induced by natural spike trains. Nature 416, 433 (2002).

43 Feldman, D. E. The spike-timing dependence of plasticity. Neuron 75, 556–571 (2012).

44 Földiák, P. Forming sparse representations by local anti-Hebbian learning. Biol. Cybern. 64, 165–170 (1990).

45 Isomura, T. & Toyoizumi, T. A Local Learning Rule for Independent Component Analysis. Sci. Rep. 6, 28073 (2016).

46 Pehlevan, C., Mohan, S. & Chklovskii, D. B. Blind nonnegative source separation using biological neural networks. Neural Comput. 29, 2925–2954 (2017).

47 Isomura, T. & Toyoizumi, T. Error-gated Hebbian rule: a local learning rule for principal and independent component analysis. Sci. Rep. 8, 1835 (2018).

48 Leugering, J. & Pipa, G. A unifying framework of synaptic and intrinsic plasticity in neural populations. Neural Comput. 30, 945–986 (2018).

49 Pawlak, V., Wickens, J. R., Kirkwood, A. & Kerr, J. N. Timing is not everything: neuromodulation opens the STDP gate. Front. Syn. Neurosci. 2, 146 (2010).

50 Frémaux, N. & Gerstner, W. Neuromodulated spike-timing-dependent plasticity, and theory of three-factor learning rules. Front. Neural Circuits 9 (2016).

51 Kuśmierz, Ł., Isomura,T. & Toyoizumi, T. Learning with three factors: modulating Hebbian plasticity with errors. Curr. Opin. Neurobiol. 46, 170–177 (2017).

52 Isomura, T., Sakai, K., Kotani, K. & Jimbo, Y. Linking neuromodulated spike-timing dependent plasticity with the free-energy principle. Neural Comput. 28, 1859–1888 (2016).

53 Jimbo, Y., Kasai, N., Torimitsu, K., Tateno, T. & Robinson, H.P. A system for MEA-based multisite stimulation. IEEE Trans. Biomed. Eng. 50, 241–248 (2003).

54 Kullback, S. & Leibler, R. A. On information and sufficiency. Ann. Math. Stat. 22, 79–86 (1951).

55 Forney, G. D. Codes on graphs: Normal realizations. IEEE Trans. Info. Theory 47, 520–548 (2001).

56 Dauwels, J. On variational message passing on factor graphs. Info. Theory, 2007. ISIT 2007. IEEE Int. Sympo., IEEE (2007).

